# Insights from Genomes and Genetic Epidemiology of SARS-CoV-2 isolates from the state of Andhra Pradesh

**DOI:** 10.1101/2021.01.22.427775

**Authors:** Pallavali Roja Rani, Mohamed Imran, J. Vijaya Lakshmi, Bani Jolly, S. Afsar, Abhinav Jain, Mohit Kumar Divakar, Panyam Suresh, Disha Sharma, Nambi Rajesh, Rahul C. Bhoyar, Dasari Ankaiah, Sanaga Shanthi Kumari, Gyan Ranjan, Valluri Anitha Lavanya, Mercy Rophina, S. Umadevi, Paras Sehgal, Avula Renuka Devi, A. Surekha, Pulala Chandra Sekhar, Rajamadugu Hymavathy, P.R. Vanaja, Vinod Scaria, Sridhar Sivasubbu

## Abstract

Coronavirus disease (COVID-19) emerged from a city in China and has now spread as a global pandemic affecting millions of individuals. The causative agent, SARS-CoV-2 is being extensively studied in terms of its genetic epidemiology using genomic approaches. Andhra Pradesh is one of the major states of India with the third-largest number of COVID-19 cases with limited understanding of its genetic epidemiology. In this study, we have sequenced 293 SARS-CoV-2 genome isolates from Andhra Pradesh with a mean coverage of 13,324X. We identified 564 high-quality SARS-CoV-2 variants, out of which 15 are novel. A total of 18 variants mapped to RT-PCR primer/probe sites, and 4 variants are known to be associated with an increase in infectivity. Phylogenetic analysis of the genomes revealed the circulating SARS-CoV-2 in Andhra Pradesh majorly clustered under the clade A2a (94%), while 6% fall under the I/A3i clade, a clade previously defined to be present in large numbers in India. To the best of our knowledge, this is the most comprehensive genetic epidemiological analysis performed for the state of Andhra Pradesh.

## Introduction

The emergence of COVID-19 as a global pandemic has necessitated approaches to understand the emergence and evolution of SARS-CoV-2. Genome sequencing has emerged as one of the widely used approaches to understand the genetic epidemiology of SARS-CoV-2^1^. The availability of the complete genome of the pathogen early in the epidemic and subsequent application of genomics on a global and unprecedented scale has provided an immense opportunity to trace the introduction, spread and genetic evolution of the SARS-CoV-2 across the globe^2^.

India is now a major country affected by COVID-19 with over 10 million people affected since the initial introduction of SARS-CoV-2 into the country in 2020 and subsequent introductions through travellers across major cities. These include states with significantly large populations and air travellers like Andhra Pradesh which has a population of 49 million people, with an estimated 1.5 to 2 million people who are part of the diaspora spread across the world.

While a number of genomes have been sequenced from different states in India^3^, there is a paucity of genomic data and genetic epidemiology of SARS-CoV-2 isolates from the state of Andhra Pradesh which motivated us to study the genomes from this state in detail. In the present study, we report a total of 293 SARS-CoV-2 genomes from the state of Andhra Pradesh. To the best of our knowledge, this is the first comprehensive report of the genetic epidemiology and evolution of SARS-CoV-2 from the state of Andhra Pradesh.

## MATERIALS AND METHODS

The study is in compliance with relevant laws and institutional guidelines and in accordance with the ethical standards of the Declaration of Helsinki and approved by Institutional Human Ethics Committee (RC.No.03/IHC/kmcknl/2020, dated 03/08/2020). The patient consent has been waived by the ethics committee. RNA samples isolated from nasopharyngeal/oropharyngeal swabs of patients from a tertiary care teaching hospital (Kurnool Medical College) were used in the study. Two protocols for RNA isolation were employed for the clinical samples. The first protocol used GenoSens (SARS-CoV-2) PCR Viral RNA extraction reagents and samples were processed as per the instructions provided by the manufacturer. The second protocol involved processing the samples using the Truprep system (Molbio Diagnostics) for COVID-19. All samples were confirmed by multiplex RT-PCR (SD Biosensor Healthcare Pvt. Ltd STANDARD M nCoV Real-Time Detection kit, Meril Diagnostics - Meril COVID-19 one-step RT-PCR Kit, LABSYSTEMS - COVIDsure Multiplex Real-time RT-PCR kit, See gene Allplex 2019-nCoVAssay detection kit) were selected for the present study.

A total of 1,43,726 samples were tested between April 21 to 5th August and 10,073 samples were identified as COVID-19 positive cases. 293 samples were considered for viral genome sequencing for the present study and were selected between the dates of 27 June to 3 August 2020. Library preparation and sequencing was performed as per the COVIDSeq protocol (Illumina, USA) as described in a previous study^4^. The samples were sequenced in technical replicates. The raw binary sequence files in bcl format were demultiplexed and converted to FASTQ files.

We followed a previously published protocol for data analysis^5^. Briefly, raw FASTQ files underwent quality control with average Phred score quality of Q30 and read length of 30 bps with adapter trimming using Trimmomatic (version 0.39)^6^. The Wuhan-Hu-1 (NC_045512.2) genome was used as the reference. Replicate files were independently aligned and merged. Genomes with ≥99% coverage and ≤5% unassigned nucleotides were processed for variant calls. The variants were annotated by ANNOVAR^7^ using custom database tables for annotating the SARS-CoV-2 genome. Filtered variants were systematically compared with other viral genomes deposited in the Global Initiative on Sharing All Influenza Data (GISAID) from India and the rest of the world. Genomes from GISAID were pairwise aligned with the reference genome (NC_045512.2) using EMBOSS^8^ and variants were called using SNP-Sites^9^. Only genomes with an alignment percentage of ≥99% and degenerate bases ≤5% were used for comparative analysis. This accounts for a total of 45,830 high-quality genome sequences submitted until 26 September 2020 (Supplementary Table 1).

Phylogenetic analysis was performed as described in a previous protocol using the genomes sequenced in this study and the dataset of 3,058 genomes from India deposited in the GISAID database (Supplementary Table 2)^2^,^10^. Briefly, the consensus sequences were aligned to the Wuhan-Hu-1 genome and the phylogenetic tree was constructed using the Augur implementation of Nextstrain^11,12^. Genomes having ambiguous collection dates or Ns >5% were excluded from the analysis. The tree was loaded in Auspice for visualisation. Lineages were assigned to the genomes using the Phylogenetic Assignment of Named Global Outbreak LINeages (PANGOLIN) package^13^.

SARS-CoV-2 genetic variants with potential impacts on functional consequences validated through experimental evidence or by computational predictions were compiled from published studies and article pre-prints. Variants filtered in this study were compared with this repository to assess their possible functional impacts. The primer/probe sequences used in the molecular detection of SARS-CoV-2 were compiled from the literature and other sources^14^. This compilation included a total of 132 primer and probe sequences. The primer and probe sequences were mapped to the Wuhan-Hu-1 reference genome using BLAST to get their genomic loci. Variants filtered in this study were mapped to these genomic loci using bespoke scripts.

## RESULTS

A 200 μL of VTM (Himedia, Mumbai) with throat swab samples were used to extract the viral RNA (SD Genomics as manufacturer suggested) from subjects with symptoms of COVID-19 infection. 10 μL of viral RNA was used for RT-PCR detection by using various kits and following the protocol specified by the manufacturer (SD Biosensor Healthcare Pvt. Ltd STANDARD M nCoV Real-Time Detection kit, Meril Diagnostics-Meril COVID-19 one-step RT-PCR Kit, LABSYSTEMS - COVIDsure Multiplex RT-PCR kit, See gene Allplex 2019-nCoV Assay detection kits). Target genes, including ORF1ab gene, RNA dependent RNA polymerase (RdRP gene), Nucleocapsid protein (N gene), and Envelope protein (E gene) were simultaneously amplified and tested during the real-time PCR assay. Confirmed SARS-CoV-2 RNA extracts with Ct values 22 to 28 were further processes for whole-genome sequencing.

A total of 293 SARS-CoV-2 genomes were sequenced which generated an average of 12.7 million raw reads. After trimming, 11.2 million reads aligned to the SARS-CoV-2 reference genome (NC_045512.2) with a mapping percentage of 97.27% and 13324X coverage (Supplementary Table 3). 276 samples having genome coverage ≥99% and ≤5% unassigned nucleotides were further processed for variant calling and consensus sequence generation.

The reference-based assembly generated a total of 615 unique genetic variants. To ensure the reliability of the variant calls only variants with read frequencies ≥50% were considered for the comparative analysis. 564 such high-quality variants, 544 of which were in protein-coding regions and 20 were in UTRs. Of the protein-coding variants, 310 variants were non-synonymous, 230 were synonymous while 4 were stopgain. This analysis also contributed to 15 novel genetic variants (Supplementary Table 4). The distribution of the variants in the genomes and their annotations are summarised in Figure 1. 145 variants were annotated as deleterious by SIFT^15^. In addition, a total of 18 genetic variants mapped to diagnostic RT-PCR primer/probe sites (Supplementary Table 5). A total of 42 and 421 variants were predicted to map to potential B and T cell epitopes respectively. 11 variants spanning potential homoplasic, hypermutable and sequencing error-prone regions of the genome were filtered. The detailed annotations of the variants are summarised in Supplementary Table 6.

**Figure 1.**
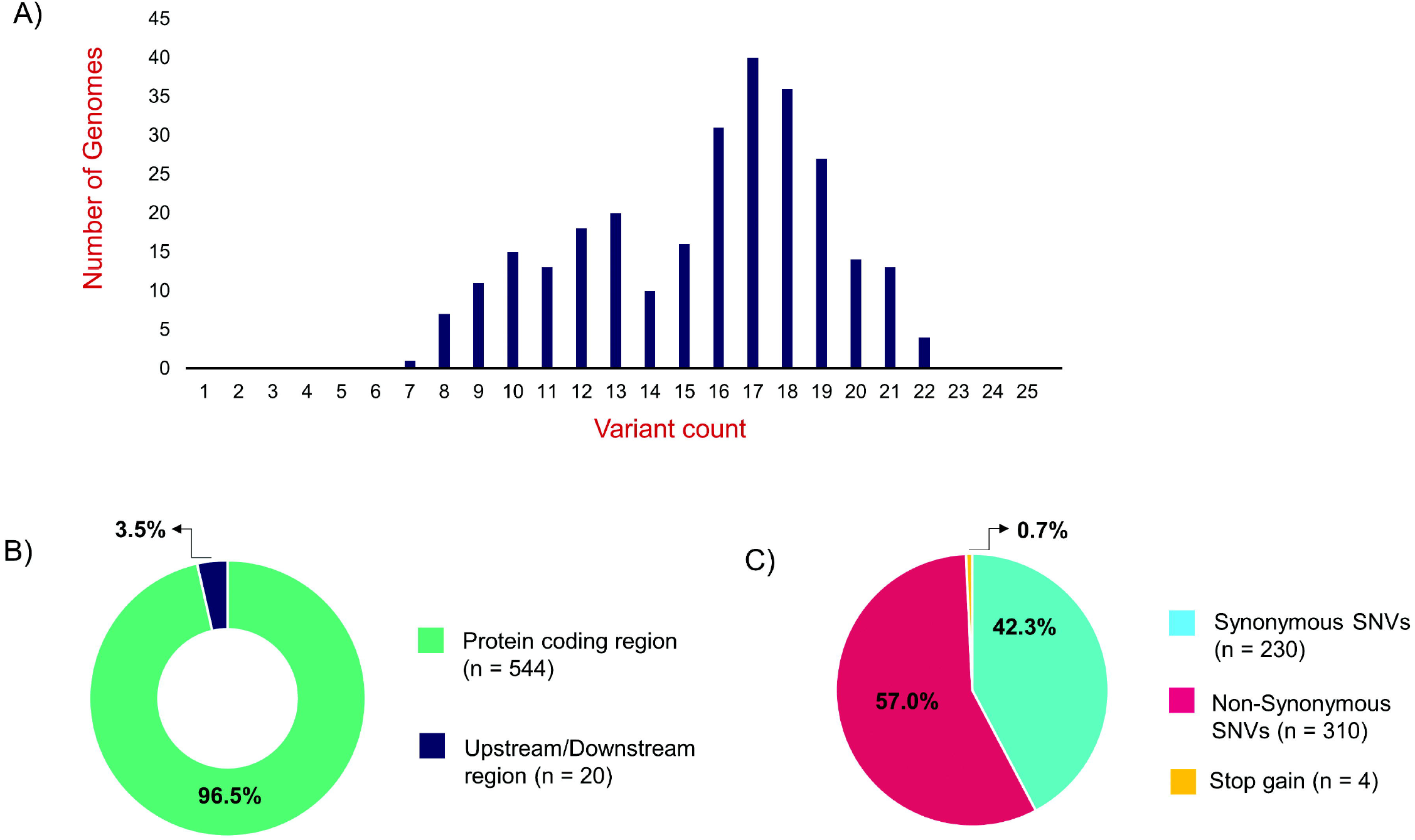
Distribution of genetic variants across genomes and their annotation summary. (A) Variant count in genomes having ≥ 99% coverage (B,C) Functional classification of the 564 variants called in this study

### Phylogenetics and unique clades and distribution

Phylogenetic analysis was done for the dataset of 3,033 SARS-CoV-2 genomes from India including 276 genomes from this study and the genome Wuhan/WH01 (EPI_ISL_406798) as the root. Out of 276 genomes, 260 genomes (94%) clustered under the clade A2a while 16 were under the clade I/A3i (6%)^16^. The phylogenetic reconstruction of the dataset of Indian genomes and the distribution of clades is summarized in Figure 2A and B. The dominant lineages for the 276 genomes, as assigned by PANGOLIN, were B.1.113 (n=129) and B.1 (n=95) as compared to other Indian genomes where B.1.1.32 and B.6 were dominant while B.1 and B.1.1 lineages were dominant for genomes in the global dataset. 5 and 1 genomes were assigned the lineages B.1.112 and B.1.104 respectively which have not been previously reported for the genomes from India (Figure 2C).

**Figure 2.**
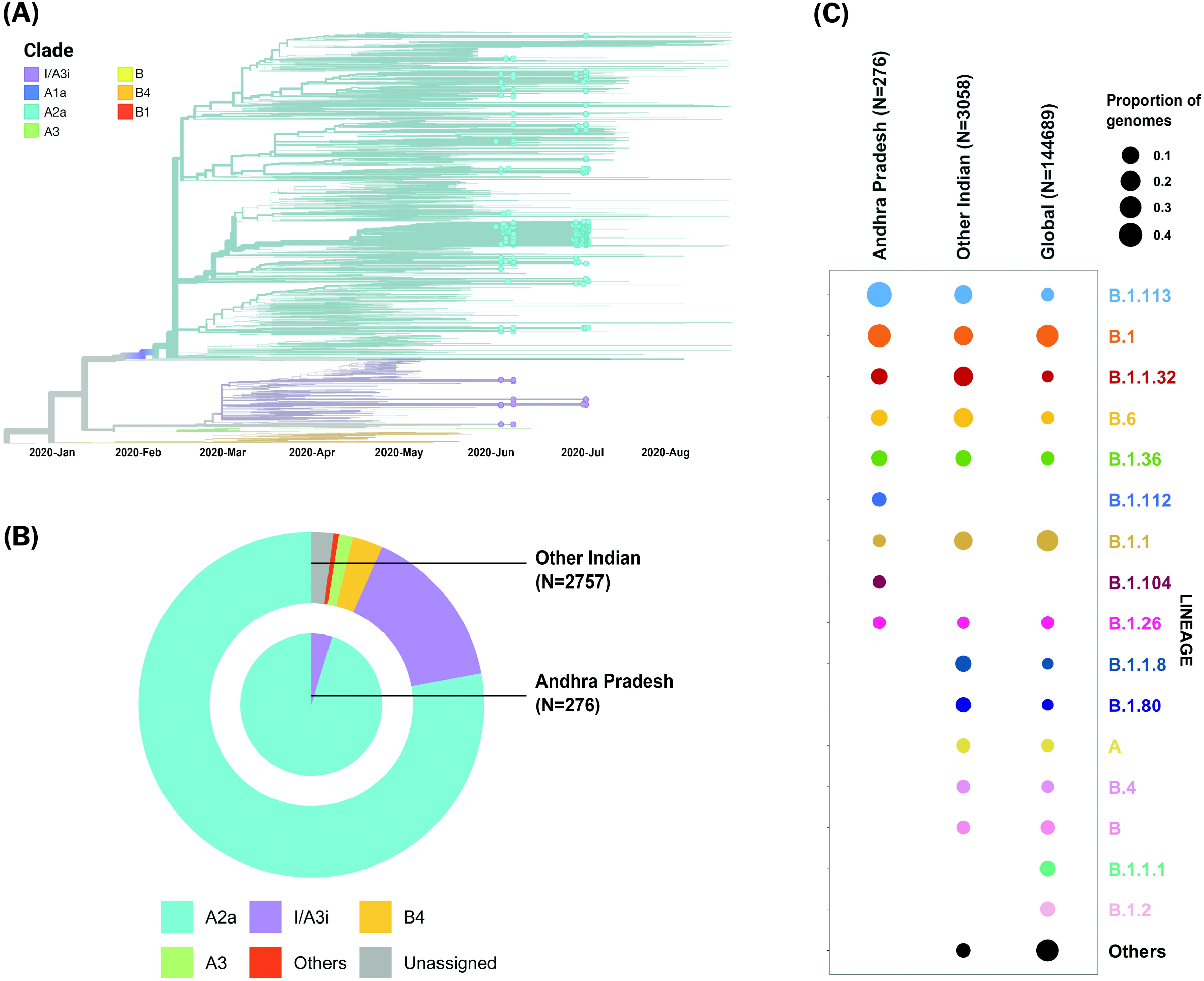
(A) Time-resolved phylogenetic reconstruction of 3033 SARS-CoV-2 genomes from India. 276 genomes from this study are highlighted (B) Distribution of clades in the 276 genomes from Andhra Pradesh and other genomes from India (C) Distribution of PANGOLIN lineages in the genomes in this study in comparison with other genomes from India and across the world.

The two earlier genomes from Andhra Pradesh (EPI_ISL_436440 and EPI_ISL_435089)^17^ sampled in the months of March and April respectively cluster under the clade I/A3i. Genomes from the neighbouring state of Telangana also show a dominant prevalence of clade I/A3i in the initial days of the pandemic and an increased representation of clade A2a in later months. The genomes sequenced in this study show a prevalence of the clade A2a in Andhra Pradesh which suggests a shift in clade dominance for this region. From the 263 genomes that cluster under clade A2a, a majority of the genomes (n=171) were seen to fall under a distinct sub-cluster that has been previously reported for genomes from Gujarat^18^. The cluster is characterized by an S194L (C28854T) mutation in the nucleocapsid protein of the virus, a mutation which was found to be significantly associated with disease mortality in Gujarat. One genome from this study (CS0804) also forms a polytomy with other samples from the neighbouring state of Telangana in the phylogenetic tree of Indian genomes, which could be suggestive of multiple, simultaneous divergence events although further data and analysis would be needed to confirm this hypothesis reliably.

### Variants associated with increased infectivity

Potential functional impact of the high-quality variants filtered in the study was identified by precisely mapping back these variants to a manually curated compilation of functionally relevant SARS-CoV-2 genetic variants. In our analysis, we were able to identify 4 genetic variants in the S gene which have been reported to be involved in increased infectivity through experimental validation. These mutations include 23403:A>G (D614G) and three co-occurring mutations 23403A>G+21575C>T (D614G+L5F), 23403A>G+24368G>T (D614G+D936Y), 23403A>G+24378C>T (D614G+S939F), having a frequency of 94.20%, 0.725%, 5.435% and 0.362% respectively in the 276 genomes analysed in this study.

## Discussion

Our analysis using the COVIDSeq approach and downstream data analysis has provided detailed insights into the genetic epidemiology and evolution of SARS-CoV-2 isolates in the state of Andhra Pradesh. A total of 564 high quality unique genetic variants were identified, out of which 15 variants are novel. Extensive analysis of the functional consequences of the filtered variants has provided insights on the impact of these genetic variants in current diagnostic practices.

Phylogenetic analysis of the genomes highlights the potential shift in clade dominance from clade I/A3i to A2a in Andhra Pradesh, a trend also observed in the neighbouring state of Telangana. The lineages B.1.112 and B.1.104 were also reported for the first time from Indian genomes.

In conclusion, our study highlights the utility of whole-genome sequencing to study the genetic landscape and evolution of SARS-CoV-2 isolates in major states like Andhra Pradesh and emphasises the use of such scalable technologies to gain better and timely insights into epidemics.

## Acknowledgements

Authors acknowledge funding from CSIR India (MLP2005). AJ, BJ, MD, and PS acknowledge research fellowships from CSIR-India. MI acknowledges a research fellowship from ICMR. DS acknowledges research fellowship from Intel India. The funders had no role in the analysis of data, preparation of the manuscript or decision to publish.

## Data Availability

The data that support the findings of this study are openly at NCBI short Read Archive with Project ID PRJNA662193 with accession IDs from SAMN16707355 to SAMN16707555. The remaining samples raw dataset are available at NCBI short Read Archive with Project ID PRJNA655577.

## Author Contributions

Sample collection, RNA isolation and RT PCR analysis: PR, JV, SA, PS, NR, DA, SK, VA, SU, AR, AS, and PC. Library Preparation and Sequencing: RCB, IM, MD, GR and PS. Assembly and variant calling DS, AJ. Analysis of Variant Functionality MR. Phylogenetics and Genetic Epidemiology BJ The manuscript was prepared by PR, AJ, RCB, BJ, MR. All authors provided critical feedback and helped to shape the analysis and manuscript. SSB and VS conceived the original idea and supervised the project.

## Supplementary Data

**Supplementary Table 1** GISAID acknowledgment table for global genomes used in the study

**Supplementary Table 2** GISAID SARS-CoV-2 genomes from India considered for phylogenetic analysis

**Supplementary Table 3** Summary of Read counts and alignment statistics of the samples used in the study

**Supplementary Table 4** Summary of novel genetic variants and their frequencies

**Supplementary Table 5** Summary of genetic variants mapping to diagnostic RT-PCR primer/probe sites.

**Supplementary Table 6** Compilation of filtered unique genetic variants along with their corresponding annotations

