## Supplementary Table 2 for "Insights from Genomes and Genetic Epidemiology of SARS-CoV-2 isolates from the state of Andhra Pradesh"

| | | | |
|---|---|---|---|
| EPI_ISL_539746 | CSIR-Centre for Cellular and Molecular Biology | CSIR-Centre for Cellular and Molecular Biology | Nikhil Hajirnis, M Soujanya Reddy, Pratheusa Maccha, Namami Gaur, Sakshi Shambhavi, Lamuk Zaveri, Shagufta Khan, Tulasi Nagabandi, Purushotham Vodnala, Payel Mukherjee, Sofia Banu, Priya Singh,Onkar Kulkarni, Dhiviya Vedagiri, Divya Gupta, Vishal Sah, Santosh Kumar Kuncha, Krishnan Harinivas Harshan, Archana Bharadwaj Siva, Karthik Bharadwaj Tallapaka,Kezia J Ann, Radhika Khandelwal, Roshan Maku Venkata, Shemin Mansuri, Sonu Uday, Rakesh K Mishra, Divya Tej Sowpati |
| EPI_ISL_539747 | CSIR-Centre for Cellular and Molecular Biology | CSIR-Centre for Cellular and Molecular Biology | Nikhil Hajirnis, M Soujanya Reddy, Pratheusa Maccha, Payel Mukherjee, Sofia Banu, Priya Singh,Onkar Kulkarni, Dhiviya Vedagiri, Divya Gupta, Vishal Sah, Santosh Kumar Kuncha, Krishnan Harinivas Harshan, Archana Bharadwaj Siva, Karthik Bharadwaj Tallapaka, Shagufta Khan, Lamuk Zaveri, Namami Gaur, Sakshi Shambhavi, Tulasi Nagabandi, Purushotham Vodnala,Deepak Kumar, Devi Prasad Vijayashankar, Disha Nanda, Divya Das, Jotin Gogoi, Manish Bhattacharjee, Rakesh K Mishra, Divya Tej Sowpati |
| EPI_ISL_539748 | CSIR-Centre for Cellular and Molecular Biology | CSIR-Centre for Cellular and Molecular Biology | Payel Mukherjee, Sofia Banu, Priya Singh, Onkar Kulkarni, Dhiviya Vedagiri, Divya Gupta, Vishal Sah, Santosh Kumar Kuncha, Krishnan Harinivas Harshan, Archana Bharadwaj Siva, Karthik Bharadwaj Tallapaka, Shagufta Khan, Lamuk Zaveri, Nikhil Hajirnis, M Soujanya Reddy, Pratheusa Maccha, Namami Gaur, Sakshi Shambhavi, Tulasi Nagabandi, Purushotham Vodnala, G. Aditya Kumar, Koushick Sivakumar, Pooja Ramesh Gupta, Rajan Kumar Jha, Shraddha Vijay Lahoti, Rakesh K Mishra, Divya Tej Sowpati |
| EPI_ISL_539749 | CSIR-Centre for Cellular and Molecular Biology | CSIR-Centre for Cellular and Molecular Biology | Payel Mukherjee, Sofia Banu, Priya Singh, Onkar Kulkarni, Dhiviya Vedagiri, Divya Gupta, Vishal Sah, Santosh Kumar Kuncha, Krishnan Harinivas Harshan, Archana Bharadwaj Siva, Karthik Bharadwaj Tallapaka, Shagufta Khan, Lamuk Zaveri, Nikhil Hajirnis, M Soujanya Reddy, Pratheusa Maccha, Namami Gaur, Sakshi Shambhavi, Tulasi Nagabandi, Purushotham Vodnala, Gokulan C G, Gunjan Purohit, Hanuman Tulashiram Kale, Pankaj Kumar, Prachand Issarapu, Rakesh K Mishra, Divya Tej Sowpati |
| EPI_ISL_539750 | CSIR-Centre for Cellular and Molecular Biology | CSIR-Centre for Cellular and Molecular Biology | Payel Mukherjee, Sofia Banu, Priya Singh, Onkar Kulkarni, Dhiviya Vedagiri, Divya Gupta, Vishal Sah, Santosh Kumar Kuncha, Krishnan Harinivas Harshan, Archana Bharadwaj Siva, Karthik Bharadwaj Tallapaka, Shagufta Khan, Lamuk Zaveri, Nikhil Hajirnis, M Soujanya Reddy, Pratheusa Maccha, Namami Gaur, Sakshi Shambhavi, Tulasi Nagabandi, Purushotham Vodnala, Rakesh K Mishra, Sonu Uday, Sudipta Mondal, Annapoorna P Karthyayani, Debabrata Jana, Debrya Saha, Divya Tej Sowpati |
| EPI_ISL_539751 | CSIR-Centre for Cellular and Molecular Biology | CSIR-Centre for Cellular and Molecular Biology | Pratheusa Maccha, Sakshi Shambhavi, Lamuk Zaveri, Shagufta Khan, Namami Gaur, Nikhil Hajirnis, M Soujanya Reddy, Tulasi Nagabandi, Purushotham Vodnala, Payel Mukherjee, Sofia Banu, Priya Singh,Onkar Kulkarni, Dhiviya Vedagiri, Divya Gupta, Vishal Sah, Santosh Kumar Kuncha, Krishnan Harinivas Harshan, Archana Bharadwaj Siva, Karthik Bharadwaj Tallapaka, G. Aditya Kumar, Koushick Sivakumar,Disha Nanda, Divya Das, Jotin Gogoi, Manish Bhattacharjee, Ravi Prasad Mukku, Rakesh K Mishra, Divya Tej Sowpati |
| EPI_ISL_539752 | CSIR-Centre for Cellular and Molecular Biology | CSIR-Centre for Cellular and Molecular Biology | Pratheusa Maccha, Shagufta Khan, Lamuk Zaveri, Namami Gaur, Sakshi Shambhavi, Tulasi Nagabandi, Nikhil Hajirnis, M Soujanya Reddy, Purushotham Vodnala, Payel Mukherjee, Sofia Banu, Priya Singh, Onkar Kulkarni, Dhiviya Vedagiri, Divya Gupta, Vishal Sah, Santosh Kumar Kuncha, Krishnan Harinivas Harshan, Archana Bharadwaj Siva, Karthik Bharadwaj Tallapaka, Disha Nanda, Divya Das, Jotin Gogoi, Manish Bhattacharjee, Ravi Prasad Mukku, Rakesh K Mishra, Divya Tej Sowpati |
| EPI_ISL_539753 | CSIR-Centre for Cellular and Molecular Biology | CSIR-Centre for Cellular and Molecular Biology | Pratheusa Maccha, Sofia Banu, Payel Mukherjee, Priya Singh, Onkar Kulkarni, Dhiviya Vedagiri, Divya Gupta, Vishal Sah, Santosh Kumar Kuncha, Krishnan Harinivas Harshan, Archana Bharadwaj Siva, Karthik Bharadwaj Tallapaka, Shagufta Khan, Lamuk Zaveri, Namami Gaur, Sakshi Shambhavi, Nikhil Hajirnis, M Soujanya Reddy, Tulasi Nagabandi, Purushotham Vodnala,Preethi Jampala, Sharada Ravi Iyer, Sulagana Mukherjee, Swetha Sundar, Peddapuvala Sai Uday Kiran, Rakesh K Mishra, Divya Tej Sowpati |
| EPI_ISL_539754 | CSIR-Centre for Cellular and Molecular Biology | CSIR-Centre for Cellular and Molecular Biology | Sakshi Shambhavi, Lamuk Zaveri, Shagufta Khan, Namami Gaur, Nikhil Hajirnis, M Soujanya Reddy, Pratheusa Maccha, Tulasi Nagabandi, Purushotham Vodnala, Payel Mukherjee, Sofia Banu, Priya Singh, Onkar Kulkarni, Dhiviya Vedagiri, Divya Gupta, Vishal Sah, Santosh Kumar Kuncha, Krishnan Harinivas Harshan, Archana Bharadwaj Siva, Karthik Bharadwaj Tallapaka, Deepak Kumar, Devi Prasad Vijayashankar, Disha Nanda, Divya Das, Jotin Gogoi, Manish Bhattacharjee, Rakesh K Mishra, Divya Tej Sowpati |
| EPI_ISL_539755 | CSIR-Centre for Cellular and Molecular Biology | CSIR-Centre for Cellular and Molecular Biology | Sakshi Shambhavi, Lamuk Zaveri, Shagufta Khan, Namami Gaur, Nikhil Hajirnis, M Soujanya Reddy, Pratheusa Maccha,Tulasi Nagabandi, Purushotham Vodnala, Payel Mukherjee, Sofia Banu, Priya Singh,Onkar Kulkarni, Dhiviya Vedagiri, Divya Gupta, Vishal Sah, Santosh Kumar Kuncha, Krishnan Harinivas Harshan, Archana Bharadwaj Siva, Karthik Bharadwaj Tallapaka, G. Aditya Kumar, Koushick Sivakumar, Pooja Ramesh Gupta, Rajan Kumar Jha, Shraddha Vijay Lahoti, Rakesh K Mishra, Divya Tej Sowpati |
| EPI_ISL_539756 | CSIR-Centre for Cellular and Molecular Biology | CSIR-Centre for Cellular and Molecular Biology | Sakshi Shambhavi, Lamuk Zaveri, Shagufta Khan, Nikhil Hajirnis, M Soujanya Reddy, Pratheusa Maccha, Namami Gaur, Tulasi Nagabandi, Purushotham Vodnala, Payel Mukherjee, Sofia Banu, Priya Singh, Onkar Kulkarni, Dhiviya Vedagiri, Divya Gupta, Vishal Sah, Santosh Kumar Kuncha, Krishnan Harinivas Harshan, Archana Bharadwaj Siva, Karthik Bharadwaj Tallapaka,G. Aditya Kumar, Koushick Sivakumar, Rakesh K Mishra, Divya Tej Sowpati |
| EPI_ISL_539757 | CSIR-Centre for Cellular and Molecular Biology | CSIR-Centre for Cellular and Molecular Biology | Shagufta Khan, Lamuk Zaveri, Namami Gaur, Sakshi Shambhavi, Nikhil Hajirnis, M Soujanya Reddy, Pratheusa Maccha, Tulasi Nagabandi, Purushotham Vodnala, Payel Mukherjee, Sofia Banu, Priya Singh, Onkar Kulkarni, Dhiviya Vedagiri, Divya Gupta, Vishal Sah, Santosh Kumar Kuncha, Krishnan Harinivas Harshan, Archana Bharadwaj Siva, Karthik Bharadwaj Tallapaka, Renu Sudhakar, Somesh Gorde, Gangumala Srinivas Reddy, Sujoy Deb, Swati Bayyana, Rakesh K Mishra, Divya Tej Sowpati |
| EPI_ISL_539758 | CSIR-Centre for Cellular and Molecular Biology | CSIR-Centre for Cellular and Molecular Biology | Shagufta Khan, Lamuk Zaveri, Namami Gaur, Sakshi Shambhavi, Nikhil Hajirnis, M Soujanya Reddy, Pratheusa Maccha, Tulasi Nagabandi, Purushotham Vodnala, Payel Mukherjee, Sofia Banu, Priya Singh, Onkar Kulkarni, Dhiviya Vedagiri, Divya Gupta, Vishal Sah, Santosh Kumar Kuncha, Krishnan Harinivas Harshan, Archana Bharadwaj Siva, Karthik Bharadwaj Tallapaka,Preethi Jampala, Sharada Ravi Iyer, Sulagana Mukherjee, Swetha Sundar, Peddapuvala Sai Uday Kiran Rakesh K Mishra, Divya Tej Sowpati |
| EPI_ISL_539759 | CSIR-Centre for Cellular and Molecular Biology | CSIR-Centre for Cellular and Molecular Biology | Shagufta Khan, Lamuk Zaveri, Namami Gaur, Sakshi Shambhavi, Nikhil Hajirnis, M Soujanya Reddy, Pratheusa Maccha,Tulasi Nagabandi, Purushotham Vodnala, Payel Mukherjee, Sofia Banu, Priya Singh, Onkar Kulkarni, Dhiviya Vedagiri, Divya Gupta, Vishal Sah, Santosh Kumar Kuncha, Krishnan Harinivas Harshan, Archana Bharadwaj Siva, Karthik Bharadwaj Tallapaka,Umesh Kumar, Unis Ahmad Bhat, Ajay Sarawagi, Priyanka Pant, Rajkanwar Nathawat, Rakesh K Mishra, Divya Tej Sowpati |
| EPI_ISL_539760 | CSIR-Centre for Cellular and Molecular Biology | CSIR-Centre for Cellular and Molecular Biology | Sofia Banu, Payel Mukherjee, Priya Singh,Onkar Kulkarni, Dhiviya Vedagiri, Divya Gupta, Vishal Sah, Santosh Kumar Kuncha, Krishnan Harinivas Harshan, Archana Bharadwaj Siva, Karthik Bharadwaj Tallapaka, Shagufta Khan, Lamuk Zaveri, Namami Gaur, Sakshi Shambhavi, Nikhil Hajirnis, M Soujanya Reddy, Pratheusa Maccha, Tulasi Nagabandi, Purushotham Vodnala, Disha Nanda, Divya Das, Jotin Gogoi, Manish Bhattacharjee, Ravi Prasad Mukku, Rakesh K Mishra, Divya Tej Sowpati |
| EPI_ISL_539761 | CSIR-Centre for Cellular and Molecular Biology | CSIR-Centre for Cellular and Molecular Biology | Sofia Banu, Payel Mukherjee, Priya Singh,Onkar Kulkarni, Dhiviya Vedagiri, Divya Gupta, Vishal Sah, Santosh Kumar Kuncha, Krishnan Harinivas Harshan, Archana Bharadwaj Siva, Karthik Bharadwaj Tallapaka, Shagufta Khan, Lamuk Zaveri, Namami Gaur, Sakshi Shambhavi, Nikhil Hajirnis, M Soujanya Reddy, Pratheusa Maccha,Tulasi Nagabandi, Purushotham Vodnala, Deepak Kumar, Devi Prasad Vijayashankar, Disha Nanda, Divya Das, Jotin Gogoi, Manish Bhattacharjee, Rakesh K Mishra, Divya Tej Sowpati |
| EPI_ISL_539762 | CSIR-Centre for Cellular and Molecular Biology | CSIR-Centre for Cellular and Molecular Biology | Sofia Banu, Payel Mukherjee, Priya Singh,Onkar Kulkarni, Dhiviya Vedagiri, Divya Gupta, Vishal Sah, Santosh Kumar Kuncha, Krishnan Harinivas Harshan, Archana Bharadwaj Siva, Karthik Bharadwaj Tallapaka, Shagufta Khan, Lamuk Zaveri, Namami Gaur, Sakshi Shambhavi, Tulasi Nagabandi, Nikhil Hajirnis, M Soujanya Reddy, Pratheusa Maccha, Purushotham Vodnala, Gokulan C G, Gunjan Purohit, Hanuman Tulashiram Kale, Pankaj Kumar, Prachand Issarapu, Rakesh K Mishra, Divya Tej Sowpati |
| EPI_ISL_539763 | CSIR-Centre for Cellular and Molecular Biology | CSIR-Centre for Cellular and Molecular Biology | Tulasi Nagabandi, Namami Gaur, Sakshi Shambhavi, Lamuk Zaveri, Shagufta Khan, Nikhil Hajirnis, M Soujanya Reddy, Pratheusa Maccha, Purushotham Vodnala, Payel Mukherjee, Sofia Banu, Priya Singh, Onkar Kulkarni, Dhiviya Vedagiri, Divya Gupta, Vishal Sah, Santosh Kumar Kuncha, Krishnan Harinivas Harshan, Archana Bharadwaj Siva, Karthik Bharadwaj Tallapaka,G. Aditya Kumar, Koushick Sivakumar, Pooja Ramesh Gupta, Rajan Kumar Jha, Shraddha Vijay Lahoti, Rakesh K Mishra, Divya Tej Sowpati |
| EPI_ISL_539764 | CSIR-Centre for Cellular and Molecular Biology | CSIR-Centre for Cellular and Molecular Biology | Tulasi Nagabandi, Namami Gaur, Sakshi Shambhavi, Lamuk Zaveri, Shagufta Khan, Nikhil Hajirnis, M Soujanya Reddy, Pratheusa Maccha, Purushotham Vodnala, Payel Mukherjee, Sofia Banu, Priya Singh,Onkar Kulkarni , Dhiviya Vedagiri, Divya Gupta, Vishal Sah, Santosh Kumar Kuncha, Krishnan Harinivas Harshan, Archana Bharadwaj Siva, Karthik Bharadwaj Tallapaka,G. Aditya Kumar, Koushick Sivakumar, Pooja Ramesh Gupta, Rajan Kumar Jha, Shraddha Vijay Lahoti, Rakesh K Mishra, Divya Tej Sowpati |
| EPI_ISL_539765 | CSIR-Centre for Cellular and Molecular Biology | CSIR-Centre for Cellular and Molecular Biology | Tulasi Nagabandi, Namami Gaur, Sakshi Shambhavi, Lamuk Zaveri, Shagufta Khan, Nikhil Hajirnis, M Soujanya Reddy, Pratheusa Maccha, Purushotham Vodnala, Payel Mukherjee, Sofia Banu, Priya Singh,Onkar Kulkarni, Dhiviya Vedagiri, Divya Gupta, Vishal Sah, Santosh Kumar Kuncha, Krishnan Harinivas Harshan, Archana Bharadwaj Siva, Karthik Bharadwaj Tallapaka,Kezia J Ann, Radhika Khandelwal, Roshan Maku Venkata, Shemin Mansuri, Sonu Uday, Rakesh K Mishra, Divya Tej Sowpati |
| EPI_ISL_539766 | CSIR-Centre for Cellular and Molecular Biology | CSIR-Centre for Cellular and Molecular Biology | Tulasi Nagabandi, Namami Gaur, Sakshi Shambhavi, Lamuk Zaveri, Shagufta Khan, Nikhil Hajirnis, M Soujanya Reddy, Pratheusa Maccha, Purushotham Vodnala, Payel Mukherjee, Sofia Banu, Priya Singh, Onkar Kulkarni, Dhiviya Vedagiri, Divya Gupta, Vishal Sah, Santosh Kumar Kuncha, Krishnan Harinivas Harshan, Archana Bharadwaj Siva, Karthik Bharadwaj Tallapaka,G. Aditya Kumar, Koushick Sivakumar, Pooja Ramesh Gupta, Rajan Kumar Jha, Shraddha Vijay Lahoti, Rakesh K Mishra, Divya Tej Sowpati |
| EPI_ISL_539767 | CSIR-Centre for Cellular and Molecular Biology | CSIR-Centre for Cellular and Molecular Biology | Tulasi Nagabandi, Namami Gaur, Sakshi Shambhavi, Lamuk Zaveri, Shagufta Khan, Nikhil Hajirnis, M Soujanya Reddy, Pratheusa Maccha, Purushotham Vodnala, Payel Mukherjee, Sofia Banu, Priya Singh,Onkar Kulkarni , Dhiviya Vedagiri, Divya Gupta, Vishal Sah, Santosh Kumar Kuncha, Krishnan Harinivas Harshan, Archana Bharadwaj Siva, Karthik Bharadwaj Tallapaka,G. Aditya Kumar, Koushick Sivakumar, Pooja Ramesh Gupta, Rajan Kumar Jha, Shraddha Vijay Lahoti, Rakesh K Mishra, Divya Tej Sowpati |
| EPI_ISL_539768 | CSIR-Centre for Cellular and Molecular Biology | CSIR-Centre for Cellular and Molecular Biology | Tulasi Nagabandi, Namami Gaur, Sakshi Shambhavi, Lamuk Zaveri, Shagufta Khan, Nikhil Hajirnis, M Soujanya Reddy, Pratheusa Maccha, Purushotham Vodnala, Payel Mukherjee, Sofia Banu, Priya Singh,Onkar Kulkarni, Dhiviya Vedagiri, Divya Gupta, Vishal Sah, Santosh Kumar Kuncha, Krishnan Harinivas Harshan, Archana Bharadwaj Siva, Karthik Bharadwaj Tallapaka,Kezia J Ann, Radhika Khandelwal, Roshan Maku Venkata, Shemin Mansuri, Sonu Uday, Rakesh K Mishra, Divya Tej Sowpati |
| EPI_ISL_539769 | CSIR-Centre for Cellular and Molecular Biology | CSIR-Centre for Cellular and Molecular Biology | Lamuk Zaveri, Shagufta Khan,Nikhil Hajirnis, M Soujanya Reddy, Pratheusa Maccha, Namami Gaur, Sakshi Shambhavi, Tulasi Nagabandi, Purushotham Vodnala, Payel Mukherjee, Sofia Banu, Priya Singh, Onkar Kulkarni, Dhiviya Vedagiri, Divya Gupta, Vishal Sah, Santosh Kumar Kuncha, Krishnan Harinivas Harshan, Archana Bharadwaj Siva, Karthik Bharadwaj Tallapaka,Umesh Kumar, Unis Ahmad Bhat, Ajay Sarawagi, Priyanka Pant, Rajkanwar Nathawat, Rakesh K Mishra, Divya Tej Sowpati |
| EPI_ISL_539770 | CSIR-Centre for Cellular and Molecular Biology | CSIR-Centre for Cellular and Molecular Biology | Lamuk Zaveri, Shagufta Khan,Nikhil Hajirnis, M Soujanya Reddy, Pratheusa Maccha, Namami Gaur, Sakshi Shambhavi, Tulasi Nagabandi, Purushotham Vodnala, Payel Mukherjee, Sofia Banu, Priya Singh, Onkar Kulkarni, Dhiviya Vedagiri, Divya Gupta, Vishal Sah, Santosh Kumar Kuncha, Krishnan Harinivas Harshan, Archana Bharadwaj Siva, Karthik Bharadwaj Tallapaka,Zeba Rizvi, Zuberwasim Sayyad, Kakade Aishwarya Arun, Amrutha H C, Ananga Ghosh, Rakesh K Mishra, Divya Tej Sowpati |
| EPI_ISL_539771 | CSIR-Centre for Cellular and Molecular Biology | CSIR-Centre for Cellular and Molecular Biology | Pratheusa Maccha, Sakshi Shambhavi, Lamuk Zaveri, Shagufta Khan, Namami Gaur, Nikhil Hajirnis, M Soujanya Reddy, Tulasi Nagabandi, Purushotham Vodnala, Payel Mukherjee, Sofia Banu, Priya Singh, Onkar Kulkarni, Dhiviya Vedagiri, Divya Gupta, Vishal Sah, Santosh Kumar Kuncha, Krishnan Harinivas Harshan, Archana Bharadwaj Siva, Karthik Bharadwaj Tallapaka, G. Aditya Kumar, Koushick Sivakumar,Disha Nanda, Divya Das, Jotin Gogoi, Manish Bhattacharjee, Ravi Prasad Mukku, Rakesh K Mishra, Divya Tej Sowpati |
| EPI_ISL_539772 | CSIR-Centre for Cellular and Molecular Biology | CSIR-Centre for Cellular and Molecular Biology | Pratheusa Maccha, Shagufta Khan, Lamuk Zaveri, Namami Gaur, Sakshi Shambhavi, Tulasi Nagabandi, Nikhil Hajirnis, M Soujanya Reddy, Purushotham Vodnala, Payel Mukherjee, Sofia Banu, Priya Singh, Onkar Kulkarni, Dhiviya Vedagiri, Divya Gupta, Vishal Sah, Santosh Kumar Kuncha, Krishnan Harinivas Harshan, Archana Bharadwaj Siva, Karthik Bharadwaj Tallapaka, Disha Nanda, Divya Das, Jotin Gogoi, Manish Bhattacharjee, Ravi Prasad Mukku, Rakesh K Mishra, Divya Tej Sowpati |
| EPI_ISL_539773 | CSIR-Centre for Cellular and Molecular Biology | CSIR-Centre for Cellular and Molecular Biology | Payel Mukherjee, Sofia Banu, Priya Singh, Onkar Kulkarni, Dhiviya Vedagiri, Divya Gupta, Vishal Sah, Santosh Kumar Kuncha, Krishnan Harinivas Harshan, Archana Bharadwaj Siva, Karthik Bharadwaj Tallapaka, Shagufta Khan, Lamuk Zaveri, Nikhil Hajirnis, M Soujanya Reddy, Pratheusa Maccha, Namami Gaur, Sakshi Shambhavi, Tulasi Nagabandi, Purushotham Vodnala, Gokulan C G, Gunjan Purohit, Hanuman Tulashiram Kale, Pankaj Kumar, Prachand Issarapu, Rakesh K Mishra, Divya Tej Sowpati |
| EPI_ISL_539774 | CSIR-Centre for Cellular and Molecular Biology | CSIR-Centre for Cellular and Molecular Biology | Payel Mukherjee, Sofia Banu, Priya Singh, Onkar Kulkarni, Dhiviya Vedagiri, Divya Gupta, Vishal Sah, Santosh Kumar Kuncha, Krishnan Harinivas Harshan, Archana Bharadwaj Siva, Karthik Bharadwaj Tallapaka, Shagufta Khan, Lamuk Zaveri, Nikhil Hajirnis, M Soujanya Reddy, Pratheusa Maccha, Namami Gaur, Sakshi Shambhavi, Tulasi Nagabandi, Purushotham Vodnala, Rakesh K Mishra, Sonu Uday, Sudipta Mondal, Annapoorna P Karthyayani, Debabrata Jana, Debrya Saha, Divya Tej Sowpati |
| EPI_ISL_539775 | CSIR-Centre for Cellular and Molecular Biology | CSIR-Centre for Cellular and Molecular Biology | Payel Mukherjee, Sofia Banu, Priya Singh, Onkar Kulkarni, Dhiviya Vedagiri, Divya Gupta, Vishal Sah, Santosh Kumar Kuncha, Krishnan Harinivas Harshan, Archana Bharadwaj Siva, Karthik Bharadwaj Tallapaka, Shagufta Khan, Lamuk Zaveri, Nikhil Hajirnis, M Soujanya Reddy, Pratheusa Maccha, Namami Gaur, Sakshi Shambhavi, Tulasi Nagabandi, Purushotham Vodnala, Gokulan C G, Gunjan Purohit, Hanuman Tulashiram Kale, Pankaj Kumar, Prachand Issarapu, Rakesh K Mishra, Divya Tej Sowpati |
| EPI_ISL_541681, EPI_ISL_541682, EPI_ISL_541683, EPI_ISL_541684, EPI_ISL_541685, EPI_ISL_541686, EPI_ISL_541687, EPI_ISL_541688, EPI_ISL_541689, EPI_ISL_541690, EPI_ISL_541691, EPI_ISL_541692, EPI_ISL_541693, EPI_ISL_541694, EPI_ISL_541695, EPI_ISL_541696, EPI_ISL_541697, EPI_ISL_541698, EPI_ISL_541699, EPI_ISL_541700, EPI_ISL_541701, EPI_ISL_541702, EPI_ISL_541703, EPI_ISL_541704, EPI_ISL_541705, EPI_ISL_541706, EPI_ISL_541707, EPI_ISL_541708, EPI_ISL_541709, EPI_ISL_541710, EPI_ISL_541711, EPI_ISL_541712, EPI_ISL_541713, EPI_ISL_541714, EPI_ISL_541715, EPI_ISL_541716, EPI_ISL_541717, EPI_ISL_541718, EPI_ISL_541719, EPI_ISL_541720, EPI_ISL_541721, EPI_ISL_541722, EPI_ISL_541723, EPI_ISL_541724, EPI_ISL_541725, EPI_ISL_541726, EPI_ISL_541727, EPI_ISL_541728, EPI_ISL_541729, EPI_ISL_541730, EPI_ISL_541731, EPI_ISL_541732, EPI_ISL_541733, EPI_ISL_541734, EPI_ISL_541735, EPI_ISL_541736, EPI_ISL_541737, EPI_ISL_541738, EPI_ISL_541739, EPI_ISL_541740, EPI_ISL_541741, EPI_ISL_541742, EPI_ISL_541743, EPI_ISL_541744, EPI_ISL_541745, EPI_ISL_541746, EPI_ISL_541747, EPI_ISL_541748, EPI_ISL_541749, EPI_ISL_541750, EPI_ISL_541751 | | | |
| see above | National Institute of Virology, NIV Influenza | National Institute of Virology, NIV Influenza | Potdar V |
| EPI_ISL_547584 | CMS, Roorkee | CSIR-Institute of Microbial Technology | Kanika Bansal, Sanjeet Kumar, Anu Singh, Debarghya Ghose, Rajesh Kumar Mishra, Dipak Dutta, Sanjeev Khosla, Prabhu B. Patil |
| EPI_ISL_547585, EPI_ISL_547586, EPI_ISL_547587, EPI_ISL_547588, EPI_ISL_547589, EPI_ISL_547590, EPI_ISL_547591, EPI_ISL_547592, EPI_ISL_547593 | Civil Hospital, Panchkula | CSIR-Institute of Microbial Technology | Kanika Bansal, Sanjeet Kumar, Anu Singh, Debarghya Ghose, Rajesh Kumar Mishra, Dipak Dutta, Sanjeev Khosla, Prabhu B. Patil |
| EPI_ISL_547870, EPI_ISL_547871, EPI_ISL_547872, EPI_ISL_547873, EPI_ISL_547874, EPI_ISL_547875, EPI_ISL_547876, EPI_ISL_547877 | Maximum Containment Laboratory, National Institute of Virology | Maximum Containment Laboratory, National Institute of Virology | Yadav,P.D., Shete-aich,A., Nyayanit,D.A., Abraham,P. |
| EPI_ISL_560318 | Civil Hospital, Panchkula | CSIR-Institute of Microbial Technology | Kanika Bansal, Sanjeet Kumar, Anu Singh, Debarghya Ghose, Amandeep Kaur, Rajesh Kumar Mishra, Poushali Chakraborty, Harsh Goar, Navin Baid, Ashwani Kumar, Dipak Dutta, Sanjeev Khosla, Prabhu B. Patil |
| EPI_ISL_560319 | IDSP unit, Dehradun | CSIR-Institute of Microbial Technology | Kanika Bansal, Sanjeet Kumar, Anu Singh, Debarghya Ghose, Amandeep Kaur, Rajesh Kumar Mishra, Poushali Chakraborty, Harsh Goar, Navin Baid, Ashwani Kumar, Dipak Dutta, Sanjeev Khosla, Prabhu B. Patil |
| EPI_ISL_560320, EPI_ISL_560321 | CMS, Roorkee | CSIR-Institute of Microbial Technology | Kanika Bansal, Sanjeet Kumar, Anu Singh, Debarghya Ghose, Amandeep Kaur, Rajesh Kumar Mishra, Poushali Chakraborty, Harsh Goar, Navin Baid, Ashwani Kumar, Dipak Dutta, Sanjeev Khosla, Prabhu B. Patil |
| EPI_ISL_560322, EPI_ISL_560323, EPI_ISL_560324, EPI_ISL_561337, EPI_ISL_561338, EPI_ISL_561339, EPI_ISL_561340, EPI_ISL_561341, EPI_ISL_561342, EPI_ISL_561343 | Civil Hospital, Panchkula | CSIR-Institute of Microbial Technology | Kanika Bansal, Sanjeet Kumar, Anu Singh, Debarghya Ghose, Rajesh Kumar Mishra, Dipak Dutta, Sanjeev Khosla, Prabhu B. Patil |
| EPI_ISL_561344 | Civil Hospital, Rupnagar | CSIR-Institute of Microbial Technology | Kanika Bansal, Sanjeet Kumar, Anu Singh, Debarghya Ghose, Amandeep Kaur, Rajesh Kumar Mishra, Poushali Chakraborty, Harsh Goar, Navin Baid, Ashwani Kumar, Dipak Dutta, Sanjeev Khosla, Prabhu B. Patil |
| EPI_ISL_569859, EPI_ISL_569860, EPI_ISL_569861, EPI_ISL_569862, EPI_ISL_569863 | CSIR-Indian Institute of Chemical Biology, MEDICA Supercpecialty Hospital Kolkata | CSIR-Indian Institute of Chemical Biology, MEDICA Supercpecialty Hospital Kolkata | Sujay Krishna Maity, Priyanka Mallick, Debaleena Bhowmik, Abhishake Lahiri, Dr. Aviral Roy, Dr. Soumen Saha, Dr. Arpita Ghosh Mitra, Dr. Rajesh Pandey, Dr. Sandip Paul, Dr. Partha Chakrabarti, Dr. Saikat Chakrabarti |
| EPI_ISL_577639, EPI_ISL_577640, EPI_ISL_577641, EPI_ISL_577642, EPI_ISL_577643, EPI_ISL_577644, EPI_ISL_577645, EPI_ISL_577646, EPI_ISL_577647, EPI_ISL_577648, EPI_ISL_577649, EPI_ISL_577650, EPI_ISL_577651, EPI_ISL_577652, EPI_ISL_577653, EPI_ISL_577654, EPI_ISL_577655, EPI_ISL_577656, EPI_ISL_577657, EPI_ISL_577658, EPI_ISL_577659, EPI_ISL_577660, EPI_ISL_577661, EPI_ISL_577662, EPI_ISL_577663, EPI_ISL_577664, EPI_ISL_577665, EPI_ISL_577666, EPI_ISL_577667, EPI_ISL_577668, EPI_ISL_577669, EPI_ISL_577670, EPI_ISL_577671, EPI_ISL_577672, EPI_ISL_577673, EPI_ISL_577674, EPI_ISL_577675, EPI_ISL_577676, EPI_ISL_577677, EPI_ISL_577678, EPI_ISL_577679, EPI_ISL_577680, EPI_ISL_577681, EPI_ISL_577682, EPI_ISL_577683, EPI_ISL_577684, EPI_ISL_577685, EPI_ISL_577686, EPI_ISL_577687, EPI_ISL_577688, EPI_ISL_577689, EPI_ISL_577690, EPI_ISL_577691, EPI_ISL_577692, EPI_ISL_577693, EPI_ISL_577694, EPI_ISL_577695, EPI_ISL_577696, EPI_ISL_577697, EPI_ISL_577698, EPI_ISL_577699, EPI_ISL_577700, EPI_ISL_577701, EPI_ISL_577702, EPI_ISL_577703, EPI_ISL_577704, EPI_ISL_577705, EPI_ISL_577706, EPI_ISL_577707, EPI_ISL_577708, EPI_ISL_577709, EPI_ISL_577710, EPI_ISL_577711, EPI_ISL_577712, EPI_ISL_577713, EPI_ISL_577714, EPI_ISL_577715, EPI_ISL_577716, EPI_ISL_577717, EPI_ISL_577718, EPI_ISL_577719, EPI_ISL_577720, EPI_ISL_577721, EPI_ISL_577722, EPI_ISL_577723, EPI_ISL_577724, EPI_ISL_577725, EPI_ISL_577726, EPI_ISL_577727, EPI_ISL_577728, EPI_ISL_577729, EPI_ISL_577730, EPI_ISL_577731, EPI_ISL_577732, EPI_ISL_577733 | | | |
| see above | NIV Influenza | NIV Influenza | Potdar V |
| EPI_ISL_578080 | CSIR-Indian Institute of Chemical Biology, MEDICA Supercpecialty Hospital Kolkata | CSIR-Indian Institute of Chemical Biology, MEDICA Supercpecialty Hospital Kolkata | Sujay Krishna Maity, Priyanka Mallick, Debaleena Bhowmik, Abhishake Lahiri, Dr. AviralRoy, Dr. Soumen Saha, Dr. Arpita Ghosh Mitra, Dr. Rajesh Pandey, Dr. Sandip Paul, Dr.Partha Chakrabarti, Dr. Saikat Chakrabarti |

| | | | |
|---|---|---|---|
| EPI_ISL_578081, EPI_ISL_578168, EPI_ISL_578175, EPI_ISL_578178, EPI_ISL_578179, EPI_ISL_578180, EPI_ISL_578181, EPI_ISL_578182, EPI_ISL_578183, EPI_ISL_578184, EPI_ISL_581449, EPI_ISL_581450, EPI_ISL_581451, EPI_ISL_581494, EPI_ISL_581495, EPI_ISL_581496, EPI_ISL_581497, EPI_ISL_581499, EPI_ISL_581500, EPI_ISL_581501, EPI_ISL_581502 | | | |
| see above | CSIR-Indian Institute of Chemical Biology, MEDICA Supercpecialty Hospital Kolkata | CSIR-Indian Institute of Chemical Biology, MEDICA Supercpecialty Hospital Kolkata | Sujay Krishna Maity, Priyanka Mallick, Debaleena Bhowmik, Abhishake Lahiri, Dr. Aviral Roy, Dr. Soumen Saha, Dr. Arpita Ghosh Mitra, Dr. Rajesh Pandey, Dr. Sandip Paul, Dr. Partha Chakrabarti, Dr. Saikat Chakrabarti |
| EPI_ISL_581503, EPI_ISL_581504 | CSIR-Indian Institute of Chemical Biology, MEDICA Supercpecialty Hospital Kolkata | CSIR-Indian Institute of Chemical Biology, MEDICA Supercpecialty Hospital Kolkata | Sujay Krishna Maity, Priyanka Mallick, Debaleena Bhowmik, Abhishake Lahiri, Dr. AviralRoy, Dr. Soumen Saha, Dr. Arpita Ghosh Mitra, Dr. Rajesh Pandey, Dr. Sandip Paul, Dr.Partha Chakrabarti, Dr. Saikat Chakrabarti |
| EPI_ISL_581505, EPI_ISL_581506, EPI_ISL_581507 | NCDC/IGIB | NCDC/IGIB | Vivekanand A, Mahesh S. Dhar, Bharathram Uppili, Nishu Tyagi, Pooja Sharma, Akshay Kanakan, Simmi Tiwari, RadhaKrishnan VS, Robin Marwal, Azka Khan, Ajit Shewale, Tushar Nale, Rajesh Pandey, Sandhya Kabra, Mohammed Faruq, Sujeet Singh, Anurag Agrawal, Partha Rakshit |
| EPI_ISL_582028 | CSIR-Indian Institute of Chemical Biology, MEDICA Supercpecialty Hospital Kolkata | CSIR-Indian Institute of Chemical Biology, MEDICA Supercpecialty Hospital Kolkata | Sujay Krishna Maity, Priyanka Mallick, Debaleena Bhowmik, Abhishake Lahiri, Dr. Aviral Roy, Dr. Soumen Saha, Dr. Arpita Ghosh Mitra, Dr. Rajesh Pandey, Dr. Sandip Paul, Dr. Partha Chakrabarti, Dr. Saikat Chakrabarti |
| EPI_ISL_582241 | NIV Influenza | NIV Influenza | Potdar V |
| EPI_ISL_583914 | DH | Department of Neurovirology, National Institute of Mental Health and Neuroscience (NIMHANS) | Chitra Pattabiraman,Vijayalakshmi Reddy, Harsha PK, Risha Rasheed, Pramada Prasad, Shafeeq S Hameed, Manjunatha Venkataswamy, Anita Desai, Ravi Vasanthapuram |
| EPI_ISL_583915 | Victoria Hospital | Department of Neurovirology, National Institute of Mental Health and Neuroscience (NIMHANS) | Chitra Pattabiraman,Vijayalakshmi Reddy, Harsha PK, Risha Rasheed, Pramada Prasad, Shafeeq S Hameed, Manjunatha Venkataswamy, Anita Desai, Ravi Vasanthapuram |
